# Motifs of the C-terminal Domain of MCM9 Direct Localization to Sites of Mitomycin-C Damage for RAD51 Recruitment

**DOI:** 10.1101/2020.07.29.227678

**Authors:** David R. McKinzey, Shivasankari Gomathinayagam, Wezley C. Griffin, Kathleen N. Klinzing, Elizabeth P. Jeffries, Aleksandar Rajkovic, Michael A. Trakselis

**Affiliations:** Department of Chemistry and Biochemistry, Baylor University, Waco, TX; Department of Chemistry, University of Pittsburgh, Pittsburgh, PA; Department of Pathology, University of California San Francisco, San Francisco, CA; Department of Obstetrics, Gynecology and Reproductive Sciences, University of California San Francisco, San Francisco, CA; Institute of Human Genetics, University of California San Francisco, San Francisco, CA; R&Q Solutions, LLC, Monroeville, PA

**Author notes:** To whom correspondence should be addressed: Michael A. Trakselis, One Bear Place #97365, Waco, TX 76798.

**Keywords:** DNA repair, homologous recombination, MCM9, NLS, Rad51, BRC variant motif, mitomycin C

## Abstract

The MCM8/9 complex is implicated in aiding fork progression and facilitating homologous recombination (HR) in response to several DNA damage agents. MCM9 itself is an outlier within the MCM family containing a long C-terminal extension (CTE) comprising 42% of the total length, but with no known functional components and high predicted disorder. In this report, we identify and characterize two unique motifs within the primarily unstructured CTE that are required for localization of MCM8/9 to sites of mitomycin C (MMC) induced DNA damage. First, an unconventional ‘bipartite-like’ nuclear localization (NLS) motif consisting of two positively charged amino acid stretches separated by a long intervening sequence is required for the nuclear import of both MCM8 and MCM9. Second, a variant of the BRC motif (BRCv), similar to that found in other HR helicases, is necessary for localization to sites of MMC damage. The MCM9-BRCv directly interacts with and recruits RAD51 downstream to MMC-induced damage to aid in DNA repair. Patient lymphocytes devoid of functional MCM9 and discrete MCM9 knockout cells have a significantly impaired ability to form RAD51 foci after MMC treatment. Therefore, the disordered CTE in MCM9 is functionally important in promoting MCM8/9 activity and in recruiting downstream interactors; thus, requiring full length MCM9 for proper DNA repair.

## Introduction

Homologous recombination (HR) of DNA involves multifaceted processes and pathways that respond to various types of DNA damage agents encountered during S/G2-phases of mitotic cells (Quinet et al. 2017; Wright et al. 2018). Recombination occurring during meiosis can generate crossovers for genetic diversity and proper segregation in germline cells, utilizing many of the same enzymes (Hunter 2015; Crickard et al. 2018). Therefore, HR is vital for genomic integrity and diversity required for organismal survival. Defects in either mitotic or meiotic HR can directly contribute to increased cancer susceptibility and infertility through improper chromosomal rearrangements that represent incomplete intermediates and are hallmarks of disease. Various DNA helicases contribute to several steps in the recombination pathways either facilitating or dissolving hybrid DNA recombinants (Huselid et al. 2020). Their individualized roles and substrate specificities in HR are commonly overlapping, making absolute distinctions of function difficult.

MCM8 and MCM9 are recent additions to the roster of DNA helicases involved in HR (Griffin et al. 2019). They are members of the ATPases associated with a variety of cellular activities (AAA+) superfamily and the minichromosome maintenance (MCM) family of proteins that includes MCM2-7 as the heterohexameric helicase complex central to the replication fork. The MCM8/9 complex does not appear to interact directly with MCM2-7, nor is it essential for replication (Gambus et al. 2013). However, MCM8/9 is commonly associated with the replication fork (Dungrawala et al. 2015) and may be able to take over helicase activities upon depletion of MCM2-7 (Natsume et al. 2017), suggesting a more active and dynamic role in elongation. Mounting evidence suggests that MCM8/9 is itself a heterohexameric complex involved in mediating unknown aspects of fork progression and/or downstream HR (Nishimura et al. 2012; Park et al. 2013; Lee et al. 2015).

Knockouts or knockdowns of MCM8 and/or MCM9 in mice and humans cause sex-specific tumorigenesis, defects in HR processing, and sensitivities to DNA damaging agents (Hartford et al. 2011; Lutzmann et al. 2012; Nishimura et al. 2012; Park et al. 2013). This results in diminished DNA damage signaling as exhibited by decreased phosphorylated CHK1 (pCHK1) and increased double-strand breaks (DSBs) as indicated by H2AX foci in the presence of various fork stalling or crosslinking agents. The absence of functional MCM8 or 9 impairs HR mediated fork rescue after damage through decreased recruitment of Mre11/Rad50/Nbs1 (MRN), RPA, and RAD51. In fact, MCM8/9 has been shown to be required for MRN nuclease activity to generate single-strand DNA (ssDNA) for HR after treatment with cisplatin (CPT) (Lee et al. 2015). Even so, there are differing reports on the temporal association of MCM8/9 in relation to RAD51 after treatment with various DNA damage agents (Nishimura et al. 2012; Park et al. 2013; Lee et al. 2015; Natsume et al. 2017). These differences may be related to differential activities of RAD51 in HR mediated fork stability/restart compared to that of direct DSB repair (Bhat et al. 2018) from specific DNA damage agents utilized, or different eukaryotic cell types.

Mutations in MCM8 and MCM9 in humans are linked to premature ovarian failure (POF) (Wood-Trageser et al. 2014; AlAsiri et al. 2015), amenorrhea, sterility (Desai et al. 2017), and cancer (Goldberg et al. 2015). In fact, deficiencies in MCM8 or 9 are phenotypically similar to Fanconi anemia (FA) patient mutations (Giampietro et al. 1993) (where ~50% of patients are infertile (Alter et al. 1991)) and suggests an overlapping role in interstrand crosslink (ICL) coupled HR, but without the associated anemia. MCM9 mutations are also linked to hereditary mixed polyposis and colorectal cancer, commonly caused by loss of function in DNA mismatch repair (MMR) (Poulogiannis et al. 2010; Traver et al. 2015). The MMR connection may be more important in regulating microhomology mediated HR that requires mismatch repair for Holliday junction progression (Spies et al. 2015; Traver et al. 2015; Tham et al. 2016) or from overlapping recognition of duplex distorting lesions (Yang 2008; Kato et al. 2017). Instead, several human cancer genomes show homo and heterozygous deletions in MCM9 coding regions, many missense mutations in MCM8 and MCM9, and altered expression levels that correlate with aggressive clinical features and poorer long-term survival in several human cancers (Kim et al. 2007; Sung et al. 2011; Lee et al. 2015; He et al. 2017). Like that found for BRCA1/2 deficient cells, MCM8 or 9 deficient cells are hypersensitive to poly(ADP-ribose) polymerase (PARP) inhibitors indicating a link between fork progression and BRAC1/2 mediated HR repair that can be exploited in the clinic for synthetically lethal therapies with platinum-based DNA crosslinking agents (Morii et al. 2019).

MCM9 contains a unique and large C-terminal extension (CTE) not found in the other MCM family members and can be alternatively spliced to give a shorter MCM9^M^ isoform that retains the conserved helicase domains but removes the CTE (Jeffries et al. 2013). The CTE is a common feature in other HR helicases and is generally considered to be unstructured with scattered putative amino acid motifs that can impact protein interactions and affect proper function (Griffin et al. 2019). However, no such motifs or role for the CTE has been identified for MCM9. Here, we can show that the CTE in MCM9 plays an essential role in nuclear import and formation of DNA repair foci after treatment with the crosslinking agent, mitomycin C (MMC). We have identified and validated a unique ‘bipartite-like’ nuclear localization signal (NLS) within the CTE that directs the nuclear import of MCM8. Finally, we have also identified a BRCv motif that is required for the recruitment of RAD51 to sites of MMC-induced damage that is analogous to that found in other HR helicases (Islam et al. 2012). The overall results confirm an influential role for the CTE of MCM9 in importing the MCM8/9 complex into the nucleus, directing it to sites of crosslink damage, and recruiting RAD51 for downstream repair.

## Results

### Specific domains of MCM9 reciprocally affect nuclear localization and foci formation after MMC damage

We and others have shown previously that MCM8 and MCM9 are generally localized to the nucleus and form nuclear foci after damage with MMC (Nishimura *et al.* 2012; AlAsiri *et al.* 2015). However, we sought to examine which of the domains of MCM9 *(**Figure 1A**)* are required for nuclear foci formation after MMC treatment. MCM9 contains a unique CTE that comprises 42% of the 1143 amino acids and 52 kDa of the total 127 kDa molecular weight of the full-length protein. The CTE is more hydrophilic compared to the rest of the protein, has a higher disorder probability, and is likely mostly unstructured (***Supplemental Figure S1***).

**Figure 1:**
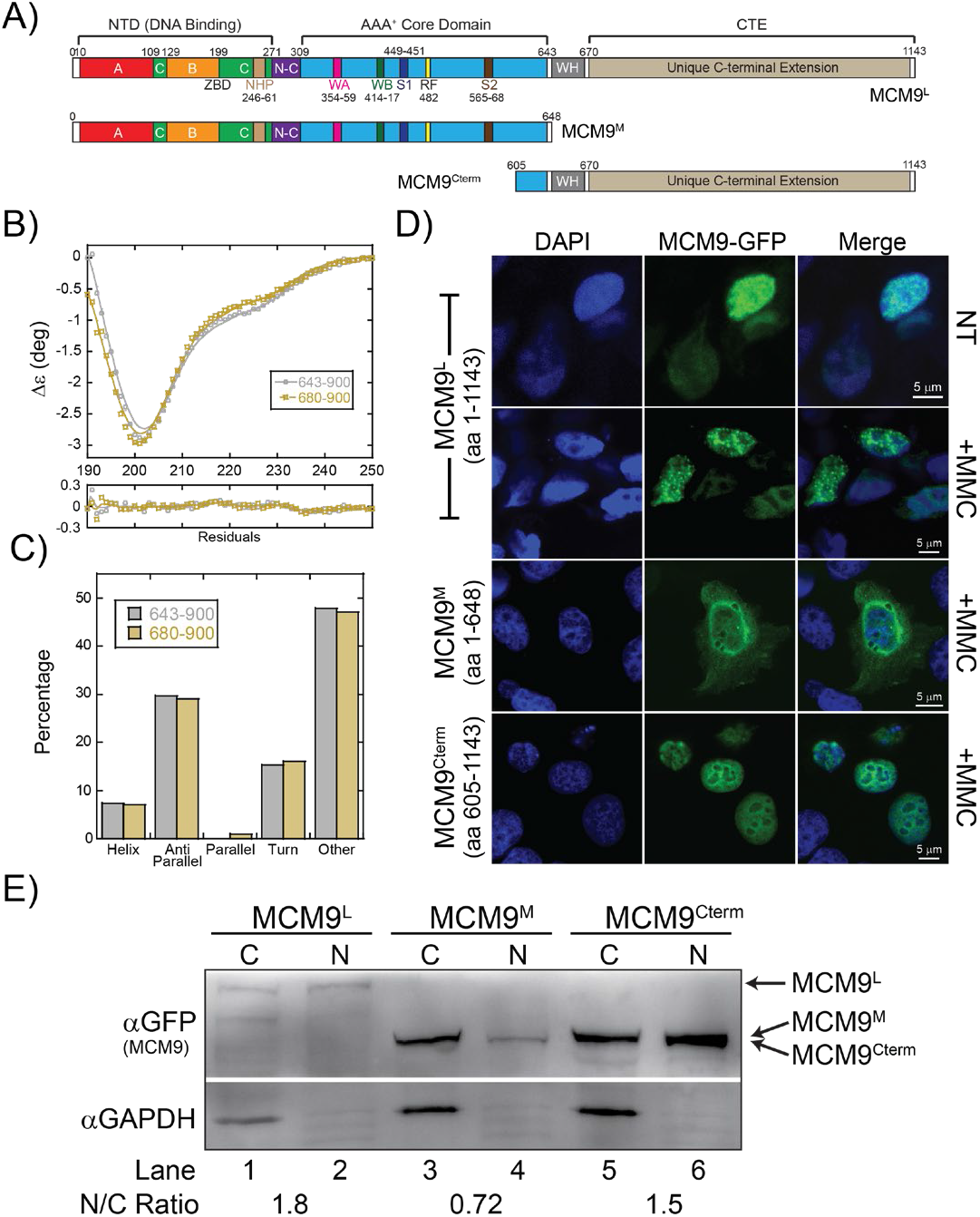
Full length MCM9 forms nuclear foci with crosslink damage directed by a disordered CTE. A) Schematic of the MCM9 linear sequence identifying known domains and motifs and separating MCM9^L^ (1-1143), MCM9^M^ (1-648), and MCM9^Cterm^ (605-1143). B) Circular dichroism of MCM9 C-terminal truncations, 643-900 (grey, open circles) or 680-900 (sand, open boxes), show a primarily unstructured CTE. Residuals of the fits (lines) to the data are shown below. C) Plot of the percentage of predicted secondary structure for each truncation based on the fit of the CD data. D) Transfection of MCM9L-GFP in HEK293T cells is nuclear and forms foci after treatment with MMC, while MCM9^M^ is primarily cytoplasmic and the MCM9^Cterm^ is fully nuclear but without MMC induced foci. E) Nuclear and cytoplasmic extractions of GFP transfected MCM9 constructs. GAPDH is used as a control for cytoplasmic proteins. Ratios of nuclear to cytoplasmic (N/C) are indicated below the image for either APB or FeBABE mapping. DNA markers (M) indicate 18 & 50 bases and fork DNA. Error bars represent standard error from 3-5 independent experiments. The products were run on a 20% native PAGE gel. p-values are defined as *< 0.05, **< 0.01 ***<0.001.

As the CTE of MCM9 is predicted to have high disorder and low secondary structure, we sought to directly measure the solution structure composition of various MCM9 CTE truncations using circular dichroism (CD). Spectra for both MCM9 643-900 and 680-900 show a pronounced minimum at 201 nm (***Figure 1B***) which is consistent with significant disorder (Chemes *et al*. 2012). There are shallow valleys from 215 to 230 nm indicative of minor β-helical and antiparallel β-sheet characteristics, but the overall spectrum is representative of a highly disordered (~50%) protein, indicated as ‘other’ in the quantification (***Figure 1C***). For the limited secondary structure, the two truncations are highly similar with only ~7% helical and ~30% β-sheet.

Full length MCM9-GFP (MCM9L) is nuclear and forms a significant number of nuclear foci upon treatment with the crosslinking agent MMC (***Figure 1D***). Interestingly, the transfection of the alternatively spliced MCM9 product (MCM9^M^) (Jeffries *et al.* 2013) showed primarily cytoplasmic staining, while MCM9^Cterm^ showed concise nuclear staining but an absence of repair foci with MMC. Nuclear and cytoplasmic extractions of various MCM9-GFP transfected constructs were used to validate these observations in a population of cells and quantified by calculating nuclear/cytoplasmic (N/C) ratios (***Figure 1E***). MCM9^L^ is more nuclear than cytoplasmic; MCM9^M^ is primarily cytoplasmic; and MCM9^Cterm^ is more nuclear, while GAPDH is used as a cytoplasmic control. Quantification of the nuclear/cytoplasmic (N/C) ratios show values above 1 for both MCM9^L^ and MCM9^Cterm^ but below 1 for MCM9^M^ that lacks the CTE. Therefore, the CTE directs MCM9 into the nucleus, but the helicase core is required for localization to DNA repair sites.

### A ‘bipartite-like’ NLS is present in the CTE of MCM9 to import MCM8/9

As MCM9^Cterm^ had predominantly nuclear staining but MCM9^M^ did not, we searched for nuclear localization sequences (NLS) in the CTE and found four conserved, high confidence putative NLS sequences (pNLS) *(**Supplemental Figure S2***). These four pNLS sequences were individually mutated in MCM9^L^-GFP, transfected into U2OS cells, and the localization was examined by confocal microscopy (***Figure 2A-B***). U2OS cells were used because of their larger nuclei that can be easily distinguished from cytoplasmic areas. Mutation of pNLS1 or pNLS2 resulted in primarily cytoplasmic staining indicating a lack of nuclear import. Alternatively, mutation of pNLS3 and pNLS4 had no noticeable effect on nuclear import. Interestingly, mutation of either pNLS1 or pNLS2 alone had a clear effect on nuclear import, indicating that a bipartite NLS1/NLS2 is required; however, the linker between NLS1 and NLS2 spans 67 a. a., much longer than a canonical bipartite NLS that generally has a 10-12 a.a. linker. Therefore, the import signal motif for MCM9 is ‘bipartite-like’ consisting of dual NLS1/NLS2 motifs connected by an extended linker region.

**Figure 2:**
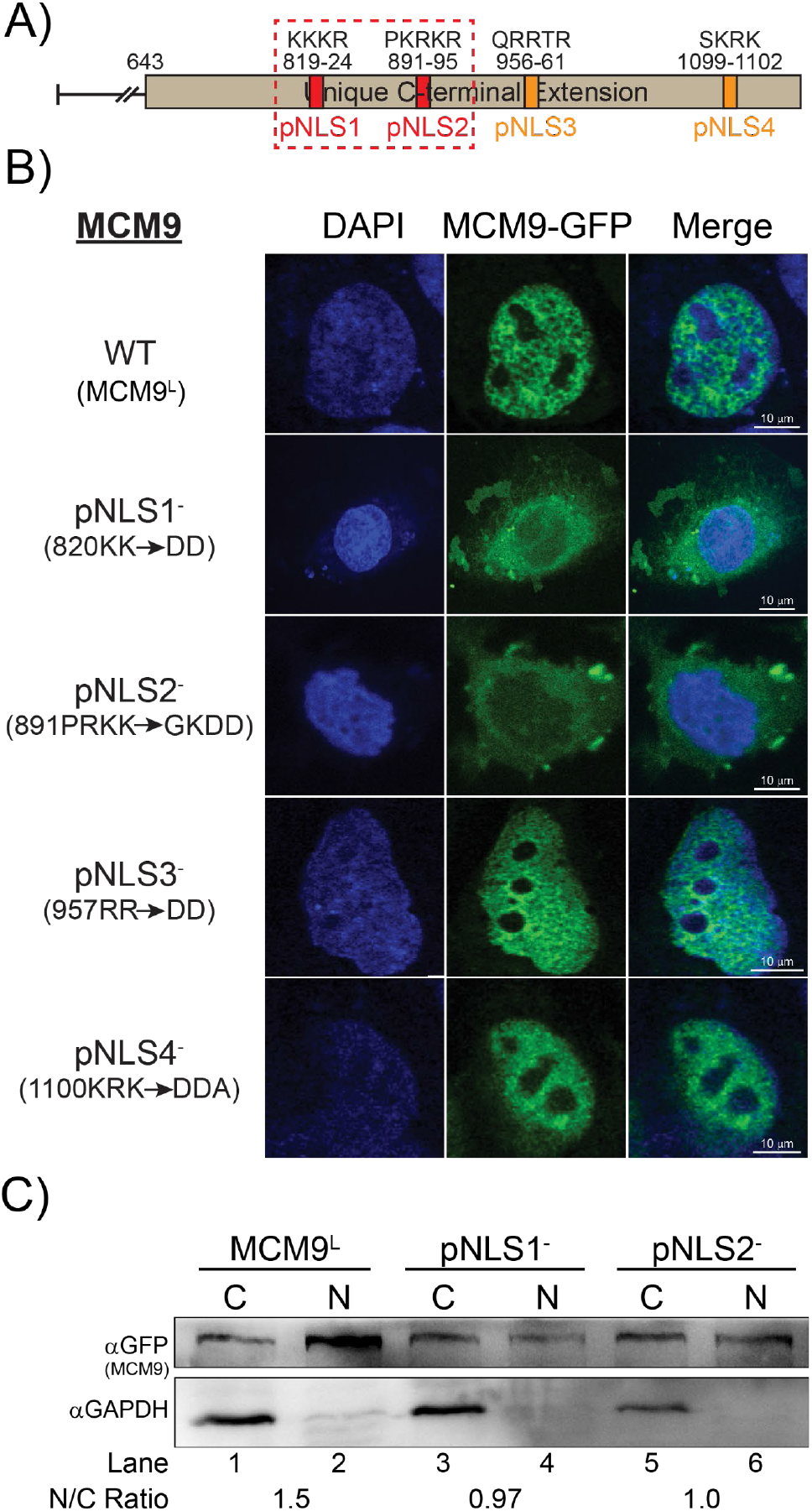
A ‘bipartite-like’ NLS1/2 is required for nuclear localization. A) Four putative nuclear localization sequences (NLS) were identified *in silico*. B) Mutation of these NLS sequences in GFP-MCM9^L^ were tested for any effect on location to the nucleus in U2OS cells. C) Nuclear and cytoplasmic extractions of GFP transfected MCM9 constructs. GAPDH is used as a control for cytoplasmic proteins. Ratios of nuclear to cytoplasmic (N/C) are indicated below the image.

To unequivocally show the effect of mutating NLS1 or NLS2 in a population of cells, we performed nuclear and cytosolic extractions on MCM9^L^-GFP construct transfected into 293T cells (***Figure 2C***). Again, WT MCM9^L^ is confined more in the nuclear fraction than the cytoplasm as indicated by N/C = 1.5. However, mutation of either NLS1 or NLS2 shows a significant decrease in the N/C ratio. This result confirms a requirement for both NLS 1 and NLS2 in the CTE of MCM9 for efficient import into the nucleus.

We also searched in silico for a pNLS in MCM8, but no high confidence motifs were detected. Therefore, we hypothesized that the NLS1/2 in the CTE of MCM9 may be responsible for nuclear import of the MCM8/9 complex. Therefore, we created a custom CRISPR-Cas9 MCM9 knockout cell line (MCM9^KO^) from 293T cells with significant multiallelic indels and validated phenotypically with severe MMC sensitivity (***Supplemental Figure S2***). To confirm if knockout of MCM9 disrupts import of MCM8 into the nucleus, MCM8-GFP was transfected into this MCM9^KO^ cell line (Figure 3). In WT 293T cells, MCM8-GFP is primarily nuclear and forms foci with MMC treatment *(**Figure 3A**),* however, in MCM9^KO^ cells, the MCM8-GFP staining is diffuse throughout the cytoplasm, not localized to the nucleus, and fails to form MMC-induced nuclear foci (***Figure 3B***). Therefore, MCM8 is imported into the nucleus complexed with MCM9 as directed by the ‘bipartite-like’ NLS1/NLS2.

**Figure 3:**
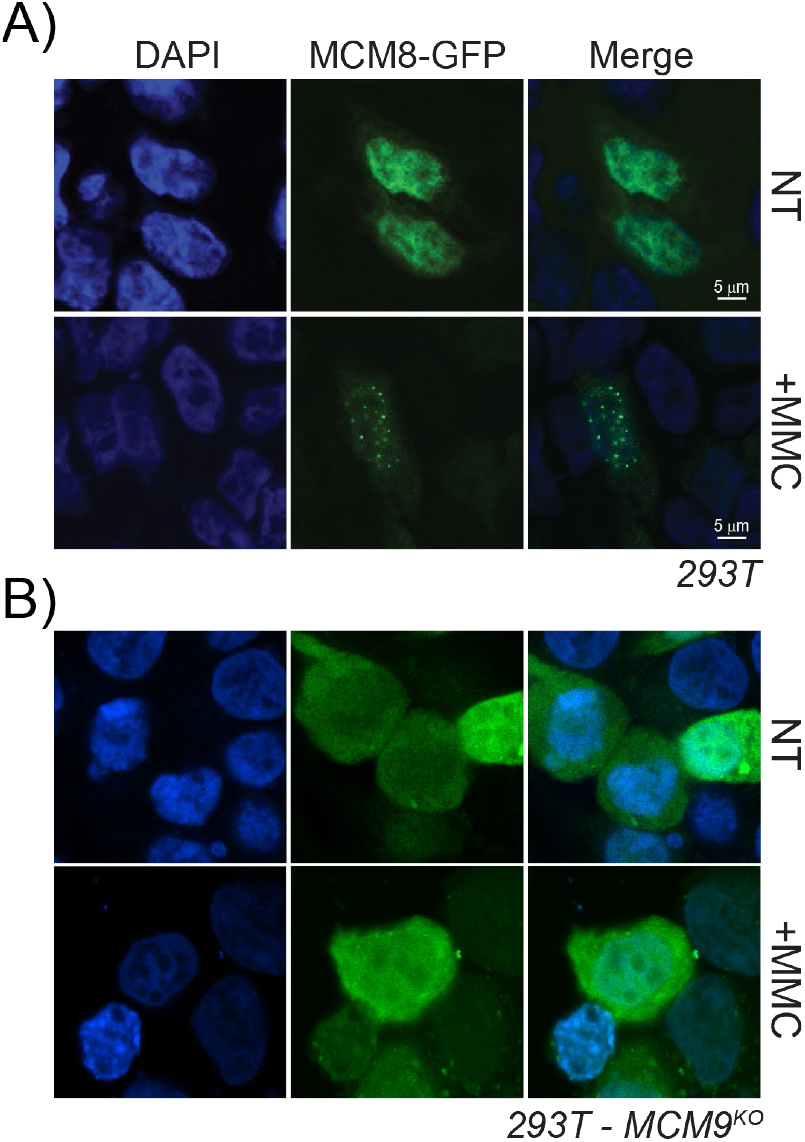
MCM9 is required for nuclear import of MCM8. MCM8-GFP was transfected into A) 293T or B) 293T-MCM9^KO^ cells in either nontreated (NT) cells or cells treated with 3 mM MMC.

### A BRCv motif is present in the CTE of MCM9

The BRC repeat sequence is a structural motif in the breast cancer type 2 susceptibility protein (BRCA2) that is utilized to promote RAD51 nucleoprotein filament formation during fork reversal or recombination (Shivji et al. 2009). The eight repeats in BRCA2 show a consensus BRC motif sequence (***Supplemental Figure S3***), and variants of these BRC motifs (BRCv) have been found in other proteins including RECQL5 helicase (Islam et al. 2012). The BRCv motif in RECQL5 is shown to be important for interacting with RAD51, directing D-loop and filament formation, and responding to crosslinking stress.

In our examination of the expansive and unique CTE of MCM9, we also identified a putative BRCv motif that is similar to that found in RECQL5 (***Figure 4A&B***). The BRCv consensus sequence within motif 1, FxTASxxxϕxϕS, is highly similar to the BRC repeats themselves FxTASGKxϕxϕS (where “ϕ” represents hydrophobic residues). BRCv motif 2 occurs after a linker region in these HR helicases that is not present in BRCA2 and is less well conserved. However, a small hydrophobic/electrostatic patch in Motif 2, ϕL(D/E)-(D/E), is present in MCM9 and similar to RECQL5. Both BRCv motifs 1 and 2 in MCM9 are well conserved across Mammalia species, but less conserved in other metazoans *(**Supplemental Figure S4B**).*

**Figure 4:**
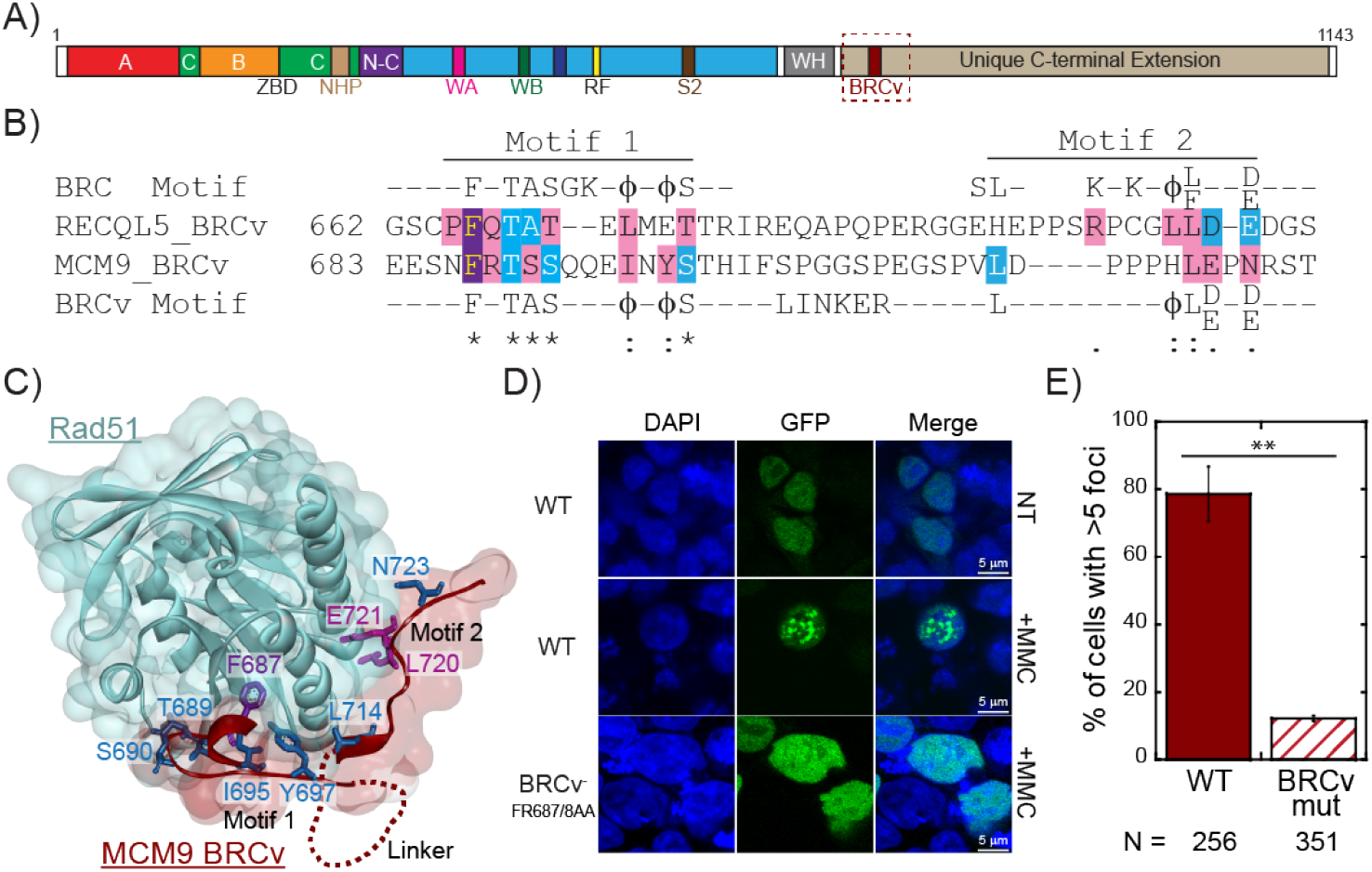
A BRCv motif in MCM9 is required for foci formation with MMC. A) A BRCv motif was identified in the CTE of MCM9 and B) aligned with the BRC consensus sequence and a previously identified BRCv motif in RecQL5. C) The MCM9 BRCv peptide was homology modelled onto the structure of RecQL5 BRCv with RAD51 (PDBID: 1N0W) visualizing the impact of conserved residues. D) Transfection of GFP-MCM9^L^(BRCv-) construct into HEK293T cells E) no longer forms significant nuclear foci with MMC treatment (** p-value < 0.01). Error bars are standard error.

Structural modeling of the MCM9 BRCv motif threaded onto BRCA2-BRC4 component in complex with RAD51 (Islam et al. 2012) show the importance of the universally conserved Phe687 residue binding deep into a hydrophobic pocket of RAD51 *(**Figure 4C*** and ***Supplemental Figure S4***). Other conserved residues (T689, S690, S691, I695, and Y697) within motif 1 make backbone and side-chain contacts across a path of hydrophobic surface area. Residues within motif 2 make both hydrophobic and hydrogen bonding contacts with RAD51 that would loop out the linker region in between *(**Supplemental Figure S4D&E**).* Leu714 is predicted to make hydrophobic contacts with RAD51 residues 254-5 and 258-9. Our model also predicts Glu722 and Asn724 makes H-bond contacts with Tyr205 and Arg250 of RAD51, respectively. Together MCM9 BRCv motifs 1 and 2 are predicted to make sufficient hydrophobic surface area contacts with RAD51 that are anchored by Phe687.

To test whether mutation of the MCM9 BRCv motif has any effect on foci formation in MMC treated cells, we transfected a mutated MCM9-GFP construct (FR687/8AA) into HEK293T cells (***Figure 4D***). The GFP fluorescence for the BRCv mutant was nuclear as expected, but MMC did not induce significant foci formation. Quantification of cells with >5 foci with MMC treatment for WT (78.7 ± 8.0%) versus BRCv^-^ (12.2 ± 0.8%) show a significant difference *(**Figure 4E**)* and illustrate the importance of the BRCv motif in directing downstream DNA repair.

### MCM9 recruitment is upstream of RAD51 during MMC treatment

As both MCM9 and RAD51 form foci during MMC treatment, we sought to examine whether there is colocalization of these two proteins during DNA damage. Previously, it was shown that the MCM8/9 complex is rapidly recruited to direct DSBs caused by *I-SceI* and promotes the association of RAD51 (Park et al. 2013; Natsume et al. 2017). Using the crosslinking agent MMC, we also observe general (but not complete) colocalization of RAD51 with MCM9 (***Figure 5A***). siRNA knockdown of RAD51 significantly reduced the RAD51 foci formation but left MCM9 foci intact *(**Figure 5A&B**).* This suggests that MCM9 is acting upstream and is required for the recruitment of RAD51 in the MMC induced DNA repair pathway.

**Figure 5:**
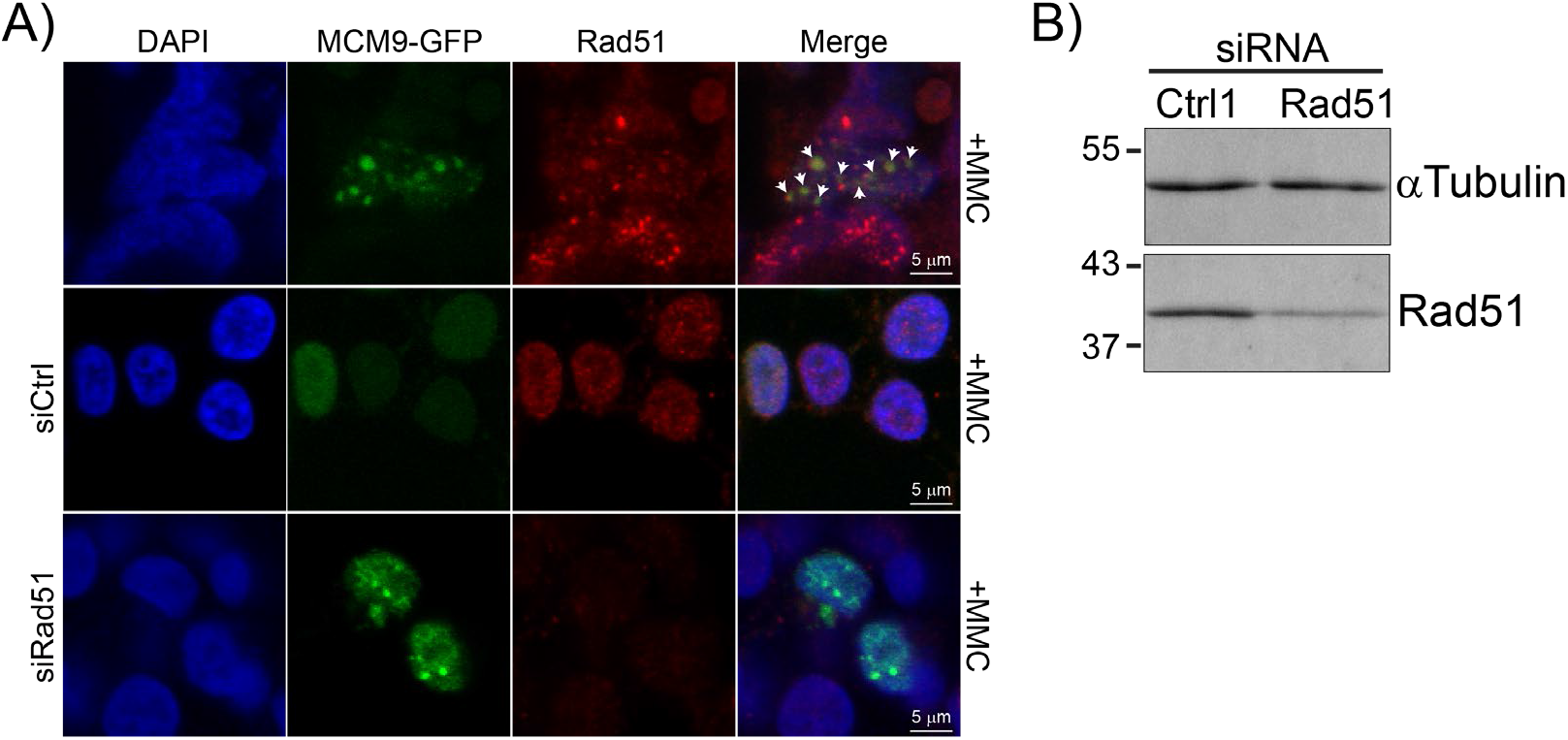
MCM9 is required for partial colocalization and recruitment of RAD51. A) MCM9 partially colocalizes with RAD51 in MMC treated HEK293T cells. White arrowheads indicate sites of colocalization. Knockdown of RAD51 B) by siRNA has no effect on MMC induced MCM9 foci formation.

As we had previously characterized MCM8 and MCM9 mutations of POF patients within consanguineous families, we examined whether these conditioned mutations in MCM9 impaired RAD51 foci formation with MMC treatment *(**Figure 6A*** and ***Supplemental Figure S5***). 8AIV-3 designates patient EBV transformed lymphocytes from an affected MCM8 family (AlAsiri et al. 2015) but is fully wild-type for both MCM8 and MCM9 and acts as a suitable control. 8AIV-3 show significant RAD51 foci after MCM damage as expected. 9BII-4 designates lymphocytes from a heterozygous family member with one wild-type allele and one splice site mutation that eliminates the CTE of MCM9 (Wood-Trageser et al. 2014). Even in these WT/MT cells, RAD51 foci formation is impaired suggesting that MCM9 copy number is important in promoting downstream repair by RAD51. Finally, 9AII-6 designates lymphocytes from a fully homozygous (MT/MT) affected patient where a nonsense mutation in exon 2 effectively creates a complete knockout of MCM9 (Wood-Trageser et al. 2014). In 9AII-6 cells, there is also a complete lack of RAD51 foci with MMC damage.

**Figure 6:**
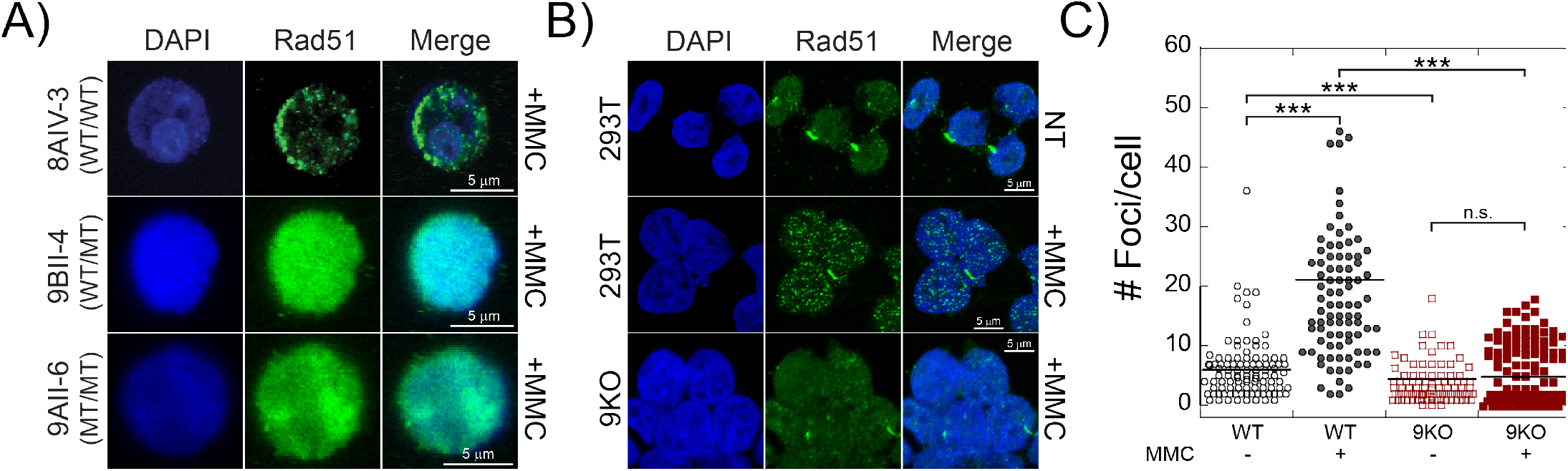
MCM9 is required for recruitment of RAD51. A) RAD51 foci after MMC treatment are disrupted in MCM9 deficient patient EBV-transformed lymphocyte cells. 8AIV-3 is wild type for both MCM8 and MCM9. 9BII-4 is heterozygous for a splice site mutation effectively eliminating the C-terminal half of MCM9. 9AII-6 is homozygous affected by a nonsense mutation in exon 2 that effectively creates a complete knockout of MCM9. B) MMC induced RAD51 foci are also C) significantly reduced in MCM9^KO^ cells and not significantly increased with MMC treatment. Foci per cell were quantified for more than 200 cells for each condition. Bar represents the average. (*** p < 0.001; n.s. not significant).

To confirm this dependency of MMC induced RAD51 foci on MCM9, we changed the experiment to quantify RAD51 foci in 293T WT cells versus their MCM9^KO^ counterparts. Treatment of 293T WT cells with MMC showed a significant increase in the number of foci per cell (6.1 ± 0.5 versus 21.2 ± 0.8), as expected *(**Figure 6B & C**).* Conversely, RAD51 foci in MCM9^KO^ cells were not significantly increased with MMC treatment (4.4 ± 0.2 versus 4.8 ± 0.2). Interestingly, the number of RAD51 foci were also significantly decreased in MCM9^KO^ cells compared to WT cells. Collectively, these data support the conclusion that RAD51 recruitment and loading for downstream repair of MMC-induced DNA damage is likely dependent on an interaction with the BRCv motif in MCM9.

### RAD51 physically interacts with the BRCv of MCM9

In order to test whether the BRCv within the CTE of MCM9 is utilized for the direct interaction with RAD51, we switched to more direct biochemical assays including a GST pulldown *(**Figure 7A***). A GST-RAD51 construct was expressed, purified, and bound to GST resin before the addition of the CTE of MCM9 (643-900 a.a.). Some MCM9 flowed through indicating either overloading or more likely a weaker interaction, but after seven washes, even more MCM9 was eluted after addition of glutathione. Further constructs of different lengths and truncations containing the BRCv mutation were prone to aggregation and insolubility issues with lower purification yields.

**Figure 7:**
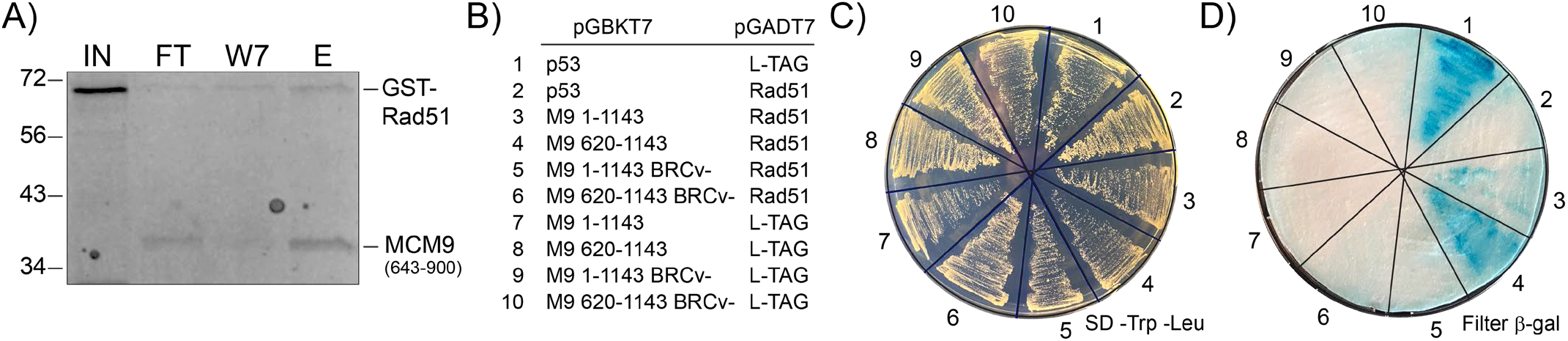
RAD51 interacts directly with the BRCv domain of MCM9. A) Pull-down of MCM9 (643-900) is present with GST-RAD51 immobilized to GST-agarose and eluted with glutathione after seven washes. IN – input GST-RAD51, FT – flow through of MCM9 (643-900), W7 – seventh wash, and E – elution. Molecular weight markers (kDa) are indicated on the left of the gel. B) Yeast two-hybrid analysis (strain: SFY526) plated on C) SD-TRP/-LEU and then D) transferred to a filter-based X-gal assay. RAD51 and MCM9 interactions confirmed through the BRCv motif of MCM9. BRCv-mutations as in ***Figure 4D***.

Therefore, we switched to a yeast two-hybrid (Y2H) assay to test the effect of a BRCv mutation on the interaction with RAD51 *(**Figure 7B-D***). Both positive (1) and negative (2, 7-10) controls grew and behaved as expected in a filter-based β-galactosidase assay: blue – positive, white – negative. Both full-length MCM9 (3) and the CTE only (4) showed blue color indicative of an interaction with RAD51. The CTE (4) was qualitatively darker blue than full length (3), but lighter than the positive control (1) possibly indicating a weaker interaction. Mutation of the BRCv motif in both constructs (5-6) eliminated the positive blue color consistent with a disruption of the RAD51 interaction.

## Discussion

The CTE domain of MCM9 is long and primarily unstructured; however, we identified two amino acid motifs that are necessary for both the import of the MCM8/9 complex and their response to crosslink damage *(**Figure 8**).* The NLS motif is ‘bipartite-like’ consisting of two positively charged amino acid stretches separated by a longer unstructured and unconventional 67 amino acid linker. The BRCv motif is utilized for localization to sites of MMC induced DNA damage and promotes the association of RAD51 for downstream repair. Interestingly, MCM9 appears to be present at the replication fork (Dungrawala et al. 2015) and can likely respond early to MMC induced damage by recruiting RAD51 for recombinational repair.

**Figure 8.**
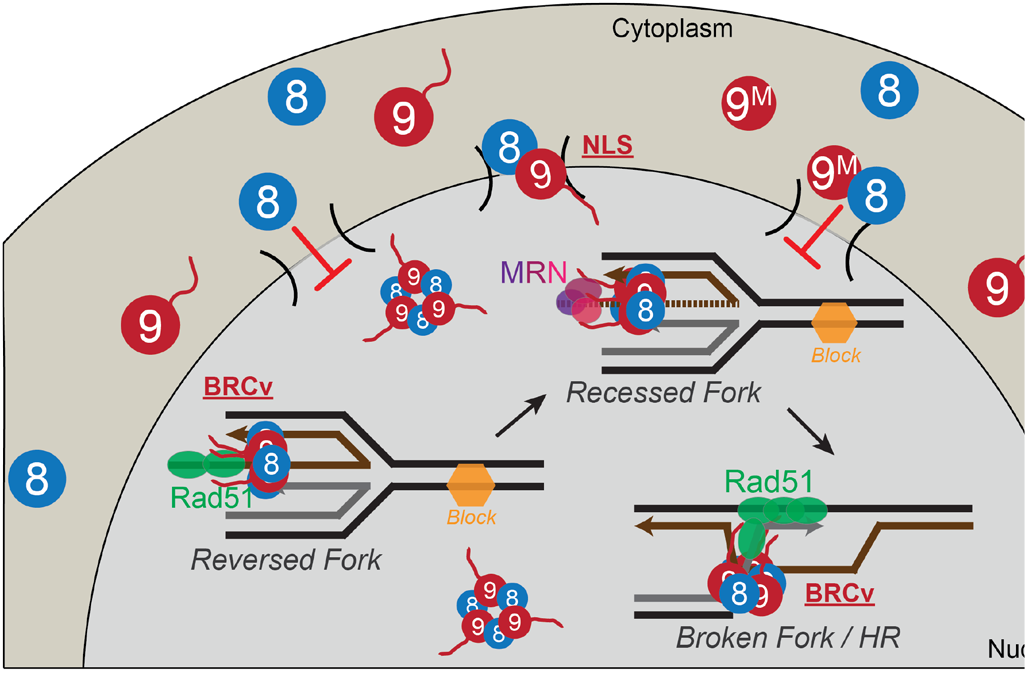
Model for MCM8/9 recruitment of RAD51 to sites of damage. The C-terminal NLS motif in MCM9 is responsible for nuclear import of MCM8/9 and the BRCv motif is required for recruiting and loading of RAD51 to sites of MMC-induced DNA damage.

### Unstructured C-terminal tails are common in DNA repair proteins

The CTE of MCM9 is translated primarily from the last exon (exon 12) and appears to be a later evolutionary addition to the MCM9 gene (Jeffries et al. 2013). MCM9 is almost universally present together with MCM8 and are distributed in most opisthokonts, excavates, and chromalveolates (Liu et al. 2009). Interestingly, MCM9 is fully present in plantae, however, the terminal exon giving rise to the CTE is absent in this eukaryotic supergroup. MCM9 members are absent in some opisthokonts, which include animals and fungi, but of those present, the primary isoform includes the CTE, while other isoforms are alternatively spliced for shorter proteins.

Several DNA replication and repair proteins contain large unstructured regions or domains that are required for proper function including several translesion synthesis DNA polymerases (Allen et al. 1991; Pryor et al. 2014), DNA replication initiation proteins (Parker et al. 2019), various DNA repair helicases (Griffin et al. 2019), mismatch repair proteins (MutLα) (Kim et al. 2019), nucleotide excision repair proteins (XPA) (Iakoucheva et al. 2001), the heterotrimeric single-stranded binding protein (RPA) (Zou et al. 2006), and even the tumor suppressor p53 (Laptenko et al. 2016). Some residues in these regions are post-translationally modified, but there are also specific motifs that are either structured on their own or adopt structure in order to control their localization, interactions, and/or functions. Previously, a role for the CTE of MCM9 was unknown. No post-translational modifications are known to occur in the CTE, but we have validated both an unconventional bipartite NLS and a BRCv motif that are required to direct function of MCM8/9 to facilitate MMC induced DNA damage repair.

### MCM9 has an unconventional ‘bipartite-like’ NLS

Classic NLS (cNLS) motifs are generally recognized as clusters of basic amino acids that are either monopartite or bipartite (Kalderon et al. 1984; Dingwall et al. 1991). Bipartite NLS motifs are conventionally described as being separated by ten to twelve residues based on the initial characterization of the cNLS of *Xenopus laveis* nucleoplasmin (Robbins et al. 1991). However, other bipartite cNLS with longer or extended linkers (20-25 residues) have been characterized in Smad4 (Xiao et al. 2003), *Aspergillus* topoisomerase II (Kim et al. 2002), and XRCC1 (Masson et al. 1998; Kirby et al. 2015), and systematic extension of the linkers in bipartite cNLS has shown that longer linkers can be utilized in nuclear import (Moore et al. 1998; Lange et al. 2010). Yet, the linker region between NLS1 and NLS2 of MCM9 is 67 amino acids and is to our knowledge one of the longest, prompting us to term this ‘bipartite-like’ NLS1/2.

Nuclear import of a bipartite NLS involves the binding of the upstream and downstream basic residues to minor and major binding pockets, respectively, for importin-α (Conti et al. 2000; Fontes et al. 2000). Both backbone and side chain interactions can be made from the linker region to a region between binding pockets. Once bound, Importin-α transports the cargo protein along with Importin-β through the nuclear pore complex depositing into the nucleus (Gorlich et al. 1995). It is hard to imagine that the longer unstructured linker between NLS1 and NLS2 in MCM9 would make specific contacts in this region, although this remains to be determined. However, we would predict that both NLS1 and NLS2 are required to bind to the binding pockets of Importin-α to facilitate import.

Based on our results, the ‘bipartite-like’ NLS1/2 in MCM9 is also responsible for the nuclear import of MCM8. MCM8 is presumed to be constitutively bound to MCM9 after translation to form a higher-order oligomeric complex. MCM8 is devoid of any identifiable NLS and based on its large molecular weight, would require an association with a partner for nuclear import. This is reminiscent of DNA ligase 3 (Lig3α) being bound to XRCC1 for co-transport into the nucleus for DNA repair (Nash et al. 1997). Another example includes the bipartite NLS in Smad4 being utilized for import of other complexed and phosphorylated Smad proteins for transcriptional regulation (Xiao et al. 2003). Interestingly, several studies have shown that knockdown or knockout of MCM8 also eliminates MCM9, but that knockdown of MCM9 has more of a singular effect (Lutzmann et al. 2012; Park et al. 2013). Therefore, we would hypothesize MCM9 is stabilized by the presence of MCM8 in an MCM8/9 complex and MCM8, although stable on its own, is unable to be imported into the nucleus. This would represent an important regulatory mechanism for ensuring that MCM8/9 is in a complex and not represented as individual components.

### MCM9 has a BRCv motif in the CTE

The BRC repeats are hallmarks within the BRCA2 protein required for interactions with RAD51 during HR and are not typically found in other proteins. A variant BRC motif (BRCv) has been characterized in the human DNA repair helicase, RECQL5, and other BRCv motifs are predicted to be present in other DNA repair helicases including yeast Srs2, Mph1, Sgs1, and Pif1 (Islam et al. 2012). The BRCv in MCM9 shows high homology to the motifs 1 [FXTA(S/T)] and 2 [ϕ(L/F)XX(D/E)] in RECQL5 with an equivalent 14 amino acid linker region between both motifs. Like that for RECQL5, BRCv motif 1 appears to be more conserved and makes more intimate contacts with the oligomerization interface of RAD51. Besides the conserved Phe687 and residues adjacent, Ile695 in MCM9 is homologous with Leu672 in RECQL5 as a small hydrophobic residue that is important for RAD51 association (Islam et al. 2012). When modeled for MCM9, Leu672 makes significant contacts in the hydrophobic patch adjacent to Phe687 making up a lynchpin for binding. Motif 2 is just adjacent to the proposed binding site for motif 1 and acts to strengthen bipartite binding to RAD51. Interestingly, the linker region is not conserved between these helicases and would represent an extruded unstructured area only required as a spacer for proper positioning of motif 2. Therefore, like RECQL5, our data support the conclusion that the BRCv motif in MCM9 is required to regulate RAD51 recruitment and filament formation to mediate recombination activities at the replication fork during times of stress.

### MCM9 aids in the recruitment of RAD51 to MMC induced sites of damage

There appears to be two conflicting viewpoints for when MCM8/9 temporally responds to various types of DNA damage in relation to RAD51. Several lines of evidence examining the cellular role of MCM8/9 come primarily from responses to the crosslinking agent, CPT, and correlate to a purported activity in later stages of HR downstream of RAD51 synapsis (Nishimura et al. 2012; Natsume et al. 2017). In those studies, RAD51 foci formed after CPT treatment are still present in MCM8 or 9 knockout cells, indicating no dependence of RAD51 on MCM8/9. This directly contradicts work here which shows no RAD51 foci in patient deficient MCM9 cells after treatment with MMC, indicating a codependence for MCM8/9 with RAD51. Instead, these results correlate with MCM8/9 acting upstream of RAD51 with MMC, consistent with other studies in human/Xenopus systems (Park et al. 2013; Lee et al. 2015). MCM8/9 are present early in the DNA repair pathway during end processing and prior to RPA binding or synapsis and colocalize with a subset of RAD51 and BRCA1 in humans (Lutzmann et al. 2012; Park et al. 2013; Lee et al. 2015). To reconcile these discrepancies, it is probable that MCM8/9 are responding to yet unknown specificities and overlapping aspects of DNA recombination associated with fork stalling and DSBs that will require further experimental depth.

Crosslinking agents such as CPT and MMC are commonly thought of in the same vein, and although both typically only cause about 5-10% ICLs, they have different targets and effects on duplex structure (Muniandy et al. 2010; Deans et al. 2011), which may be responsible for different temporal recruitment of MCM8/9. CPT crosslinking occurs at N7 of guanine at either 5’-GpG or 5’-ApG resulting primarily in <u>intra</u>strand crosslinks and <8% ICLs (Eastman 1986). MMC reacts with the N^2^ exocyclic amine of guanine preferentially at 5’-CpG sites forming monoaducts that insert into the minor groove and can form ICLs across in about 10% of products (Teng et al. 1989; Bizanek et al. 1992; Muniandy et al. 2010). However, the major difference between CPT and MMC is their effects on duplex structure. CPT induces major helical distortion with both monoadducts and ICLs underwinding the helix, bending of the strands inward, and flipping out of the unpaired cytosines (Huang et al. 1995). On the other hand, MMC causes minimal distortion, no bending, and maintains the C-G base pairs even with the G-mito-G ICL (Rink et al. 1996). These differences will be important in understanding MCM8/9 specificities, recruitment, and effects on RAD51 pathways moving forward.

Single monoaducts or intrastrand crosslinks on either the leading or lagging strands are blocks to DNA synthesis decoupling the replisome, while ICLs are total replisome blocking adducts. In these scenarios various mechanisms of recombination would be utilized including fork reversal, template switching, and single-strand breaks repaired by sister chromatin recombination. It is probable that CPT and MMC direct MCM8/9 to act differentially in recruiting RAD51 in multiple pathways for repair. Damage from CPT is primarily recognized and repaired by nucleotide excision repair (NER) (Furuta et al. 2002; Welsh et al. 2004), but because the duplex is distorted, the mismatch repair (MMR) pathway also contributes (Fink et al. 1996; Zhao et al. 2009). On the other hand, MMC lesions may only be recognized during active DNA replication processes and repaired by various translesion synthesis (TLS), HR, and NER pathways (Lee et al. 2006).

Therefore, our work here has provided evidence that the CTE within MCM9 is important in regulating MCM8/9 activities during DNA damage. Entry into the nucleus after translation is directed through an unconventional ‘bipartite-like’ NLS in the CTE of MCM9 that imports MCM8/9 as a complex. Once there, MCM8/9 associates with the replisome to actively respond to DNA damage encountered during synthesis. Recruitment of RAD51 occurs through a BRCv motif in the CTE of MCM9 that carries out downstream recombination. Even so, further efforts are needed to better understand the specific enzymatic functions and interactions of MCM8/9 in relation to different DNA damage agents to more specifically describe its reported helicase function in relation to the many other analogous HR helicases.

## Materials and Methods

### Materials

Oligonucleotides were purchased from Sigma (St. Louis, MO) or IDT (Coralville, IA). ON-TARGET *plus* siRNA to RAD51 from Dharmacon (Lafayette, CO). Mission siRNA and universal negative control #1 was from Sigma-Aldrich (St. Louis, MO). Mitomycin C (MMC) was from Thermo Fisher (Waltham, MA). Restriction enzymes from New England Biolabs (Ipswitch, MA). All other chemicals were analytical grade or better.

### Cloning

pEGFPC2-MCM9 full length has been described previously (Wood-Trageser *et al.* 2014). The truncated mutants – MCM9^Cterm^ (a.a. 605-1143), MCM9^M^ (a.a. 1-648) were created by traditional restriction site cloning into pEGFP-C2 using *XhoI/XmaI.* Codon optimized MCM9 for bacterial expression (a.a. 643-900 or 680-900) (Genewiz, South Plainfield, NJ) were cloned into pGEX-6P1 using *BamHI* and *XhoI* restriction sites. Mutants at putative nuclear localization sites: pNLS1 (820KK➔DD), NLS2 (891 PKRK➔GKDD), NLS3 (957KK➔DD), NLS4 (1099KRK >DDA), and BRCv (687FR➔AA) sites were created by Quikchange mutagenesis (Agilent Genomics, Santa Clara, CA) using Kapa DNA polymerase (Kapa Biosystems, Wilmington, MA) and screened with novel silent restriction sites. All primers are listed in ***Supplemental Table S1***. All mutations were confirmed at the Genomic Sequencing and Analysis Facility (University of Texas Austin).

### Sequence Analysis, Alignments, and Protein Homology Modelling

The amino acid sequence of human MCM9 (Accession: NP_060166.2) was analyzed using ProtScale (https://web.expasy.org/protscale/) according to the Kyte and Doolittle model of hydropathicity (Kyte *et al.* 1982). The disorder probability was calculated using DISOPRED (http://bioinf.cs.ucl.ac.uk/disopred/) (Ward *et al.* 2004). The secondary structure prediction was calculated using PSIPRED 4.0 (http://bioinf.cs.ucl.ac.uk/psipred/) (Jones 1999). The four highest confidence NLS sites for MCM9 were predicted and determined by NLS Mapper (http://nls-mapper.iab.keio.ac.jp) (Kosugi *et al.* 2009). The BRCA2 BRC4-RAD51 crystal structure (PDBID: 1N0W) was used to create a homology model for the BRCv motif of MCM9 (a.a. 682-700, 713-726) using Swiss-Model server (https://swissmodel.expasy.org/) (Waterhouse *et al.* 2018) and validated using QMEAN (https://swissmodel.expasy.org/qmean/) (Benkert *et al.* 2009).

### Cell Culture

HEK293T or U2OS cells were grown in Dulbecco’s modified Eagle’s medium (DMEM) (Corning Cellgro, Manassas, VA) with 10% fetal bovine serum (FBS) (Atlanta Biologicals, Atlanta, GA) and 1% penicillin/streptomycin (P/S) (Gibco, Gaithersburg, MD) in 5% CO_2_ at 37 °C. Patient EBV transfected lymphocytes (Wood-Trageser *et al.* 2014; AlAsiri *et al.* 2015) were grown in suspension in RPMI 1640 (Corning, Corning, NY) with 10% FBS and 1% P/S.

### Cell Transfection and Treatment

HEK293T or U2OS cells plated on poly-lysine coated coverslips were transfected with the respective EGFP-MCM9 constructs using linear polyethyleneimine (LPEI) or lipofectamine 3000 (ThermoFisher, Waltham, MA) according to manufacturer’s directions. Briefly, transfection reagent and plasmids were incubated at RT for 10 minutes and added to cells in Opti-MEM serum free media (ThermoFisher, Waltham, MA) or DMEM 10% FBS with no antibiotics. siRNA RAD51 knockdown or the universal negative control #1 were transfected alone or in combination with a plasmid using Dharmafect1 or DharmafectDuo (Dharmacon, Lafayette, CO). The transfection mixture was removed after 6 hours and replaced with DMEM / 10% FCS for up to 24 hours. The cells were then treated with indicated concentrations of MMC (Sigma-Aldrich, St. Louis, MO) for 6 hours. Post treatment, the cells were either immediately prepped for western blotting, direct confocal microscopy, or immunofluorescence. Western blots were probed with RAD51 (H-92 or sc-8349) and α-tubulin (TU-02) antibodies (Santa Cruz Biotechnology Inc., Dallas, TX). Fluorescent secondary antibodies were α-rabbit IgG-Alexa488 or α-mouse IgG-Alexa647 conjugates (ThermoFisher, Waltham, MA). Membranes were scanned using the GE LAS 4000 imager (GE Healthscience, Marlborough, MA).

### Immunofluorescence

Adherent cells were washed in PBS (2 times), fixed in 4% paraformaldehyde in PBS for 10 minutes and permeabilized with 0.1% Triton X-100 in PBS (PBST) for 15 minutes. Cells were blocked overnight with 5% BSA in PBST at 4 °C and then incubated with 1:50 RAD51 primary antibody (sc-8349, Santa Cruz or ab63801, Abcam) in 2.5% BSA in PBST for 1 hour at 37 °C. Cells were washed three times in PBST and incubated with 1:50 dilution of the appropriate fluorescent secondary antibody followed by wash with PBST (3 times). Cells were mounted in DAPI mountant (Prolong Gold, Thermo Fisher) and sealed with clear polish and imaged under a FV-1000 epifluorescence or FV-3000 confocal laser scanning microscope (Olympus Corp.). Images were processed with vendor included Fluoview (v.4.2b) or CellSens (v2.2) software. For suspension EBV transfected lymphocytes, cells were immunostained similar to adherent cells but were harvested by centrifugation after MMC treatment and immunostaining was performed in suspension. To mount the cells, they were re-suspended in one microliter of mountant and placed on a coverslip, which was then picked up using a microscopic slide, and sealed with clear polish. RAD51 foci from epifluorescence images were automatically counted from individually gated cells using identical thresholds that eliminated background noise using Image J (Rasband 1997-2016, 17 October 2015). Foci per cell are presented in a dot plot, averaged, and the standard error reported.

### Subcellular Fractionation

HEK293T cells transfected with EGFP constructs containing MCM9 truncations/mutations were cultured in 6-well plates and harvested. Fractionation into nuclear and cytoplasmic portions was carried out according to manufacturer’s directions using a Nuclear/Cytosol Fractionation Kit (K266, BioVision, Milpitas, CA). Equal volumes of protein were loaded onto a 12% acrylamide gel and transferred to PVDF membranes. Membranes were probed with the following primary antibodies: anti-GFP (Invitrogen, MA5-15256) and anti-GAPDH (DSHB, hGAPDH-2G7). Membranes were incubated with goat anti-mouse HRP conjugated secondary antibodies (Novex, A16072) and visualized using luminol reagent (Santa Cruz Biotechnology, sc-2048). Membranes were scanned using the GE LAS 4000 imager (GE Healthscience, Marlborough, MA).

### Protein Purifications

GST-tagged RAD51 was purified as described previously (Davies *et al.* 2007). C-terminally truncated MCM9 constructs (643-900 or 680-900) with GST at the N-terminus were transformed into C43 pLysS (Lucigen, Middleton, WI) and induced with 0.25 mM IPTG at OD_600_ ~ 0.5 for 4 hours at 37 °C. Pellets were resuspended in lysis buffer (20 mM HEPES pH 8.0, 300 mM NaCl, 10 mM BME, 10 mg/mL lysozyme, 0.1% Triton-X 100, 1 mM PMSF, 1 mM EDTA, 4 μg/uL pepstatin A) on ice. The clarified lysate was loaded onto the GSTrap column in Buffer A (20mM HEPES pH 8.0, 300 mM NaCl, 10 mM BME, 1 mM EDTA, 10% glycerol) and eluted with Buffer B (Buffer A + 10 mM glutathione). Fractions were concentrated using a spin concentrator and rocked overnight at 4 °C with TEV protease. The cleaved product was applied again to a GSTrap column, and the flow through collected. Purity was confirmed to be greater than 95% by SDS-PAGE, and the concentration was determined by absorbance using ε = 12,615 M^-1^cm^-1^.

### Circular Dichroism

CD spectra of 5 μM purified MCM9 643-900 and 680-900 were collected using a Jasco J-815 spectropolarimeter at room temperature (23 °C) in 10 mM potassium phosphate buffer (pH 7.0) containing 100 mM (NH_4_)_2_SO_4_ using a 0.1 cm quartz cell. Five individual spectra were collected using a 1 nm bandwidth at 0.5 nm intervals at a speed of 50 nm/min before averaging. The raw data in measured ellipticity (ε) was then analyzed using BeStSel (http://bestsel.elte.hu/index.php) for conversion to △ε units and determination of secondary structure (Micsonai *et al.* 2015; Micsonai *et al.* 2018). All data points analyzed were collected under a HT measurement of 800 V. Secondary structure predictions had NRMSD values <0.05.

### Affinity Pull-down

GST tagged RAD51 (10 μM) was added to glutathione agarose (UBPBio, Aurora, CO) at room temperature for 10 minutes before spinning down and removing the supernatant. Then, an equal molar amount of purified His-MCM9 (643-900) was added and incubated for 10 minutes. Samples were spun down and supernatant was reserved as the flow-through. The GST-agarose was washed with an equal volume of GST binding buffer (10 mM Na_2_PO_4_, 1.8 mM K_2_HPO_4_, 140 mM NaCl, and 2.7 mM KCl pH 7.3), and centrifuged for 10 seconds at 6000 g at least seven times. Proteins were eluted with an equal amount of GST elution buffer (50 mM Tris-HCl, pH 8.0, and 10 mM reduced glutathione). Eluted samples were separated using 10% SDS-PAGE gels and stained with Coomassie.

### Yeast Two-hybrid Assay

To generate the yeast two-hybrid plasmids, full-length (1-1143) or C-term (620-1143) was inserted into pGBKT7 (Clontech, Mountain View, CA) using SmaI/SalI restrictions sites. The BRCv (687FR➔AA) mutation was created by Quikchange mutagenesis (Agilent Genomics, Santa Clara, CA) using Kapa DNA polymerase (Kapa Biosystems, Wilmington, MA) and screened with novel silent restriction sites. RAD51 was PCR amplified from pGAT3-RAD51 (Davies et al. 2007) with primers that included *MfeI* and *XhoI* sites and then ligated into pGADT7 (Clontech, Mountain View, CA) digested with *EcoRI/XhoI.* The yeast strain SFY526 (MATa, ura3-52, his3-200, ade2-101, lys2-801, trp1-901, leu2-3, 112, canr, gal4-542, gal80-538, URA::GAL1 UAS-GAL1TATA-lacZ) was transformed with the appropriate plasmids (see figure legends) according to the manufacturer’s instructions using the lithium acetate procedure (Clontech Matchmaker 2 manual). Liquid cultures were grown overnight in standard dropout (SD) media lacking tryptophan and leucine. The cells were restreaked onto SD/-Trp/-Leu and incubated at 30 °C for 2–3 days before performing a colony-lift β-galactosidase assay (X-gal, Sigma-Aldrich) according to the Yeast Protocols Handbook (PT3024-1, Clontech).

## Supplemental Data

Supplementary Data are available online.

## Acknowledgments

We acknowledge the Baylor Molecular Bioscience Center (MBC) and the Center for Microscopy and Imaging (CMI) for providing instrumentation and resources aiding this project.

## Author Contributions

D.R.M. performed the cloning for the NLS mutants and associated microscopy and extraction western blots, created the CRISPR/Cas9 knockout strains, and performed the circular dichroism experiments. S.G performed and quantified the immunofluorescence on the patient lymphocytes and for the BRCv mutants. W.C.G. and K.N.K. did the cloning, protein expression, and pull-down assays for the MCM9 C-terminal truncations. E.P.J. did the cloning, transfections, and imaging of the MCM9 domain constructs. A.R. provided the patient lymphocytes through an MTA00001048 to Baylor University. M.A.T. designed the experimental approach, performed the yeast two-hybrid experiments, RAD51 foci formation, and wrote the paper. All authors analyzed data, prepared figures, and edited the manuscript.

## Abbreviations

BER: base excision repair;
BRC: breast cancer;
BRCA1: breast cancer type 1 susceptibility protein;
BRCA2: breast cancer type 2 susceptibility protein;
BME: β-mercaptoethanol;
BRCv: BRC variant motif;
BSA: bovine serum albumin;
CD: circular dichroism;
cNLS: classic NLS;
CTE: C-terminal extension;
CPT: cisplatin;
DAPI: 4’,6-diamindino-2-phenylindole;
DMEM: Dulbecco’s modified Eagle’s medium;
DSB: double-strand break;
dsDNA: double-stranded DNA;
EBV: Epstein-Barr virus;
EDTA: ethylenediaminetetraacetic acid;
FA: Fanconi anemia;
FBS: fetal bovine serum;
GAPDH: glyceraldehyde-3-phosphate dehydrogenase;
GFP: green fluorescent protein;
GST: glutathione S-transferase;
HEK: human embryonic kidney (cells);
HEPES: hydroxyethyl piperazineethanesulfonic acid;
HR: homologous recombination;
HRP: horseradish peroxidase;
ICL: interstrand crosslink;
IPTG: isopropyl β-D-thiogalactopyranosidase;
KD: knockdown;
KO: knockout;
LPEI: linear polyethyleneimine;
MCM: minichromosomal maintenance;
MMC: mitomycin-C;
MMR: mismatch repair;
MRN: Mre11/ Rad50/ Nbs1 complex;
NER: nucleotide excision repair;
NLS: nuclear localization signal;
NTD: N-terminal domain;
OD: ocular density;
PARP: poly(ADP-ribose) polymerase;
PBS: phosphate-buffered saline;
pCHK1: phosphorylated checkpoint kinase 1;
PMSF: phenylmethylsulfonyl fluoride;
pNLS: putative NLS;
PVDF: polyvinylidene difluoride;
RPA: replication protein A;
RPMI: Roswell Park Memorial Institute media;
ssDNA: single-stranded DNA;
SDS-PAGE: sodium dodecyl sulfate polyacrylamide gel electrophoresis;
TEV: tobacco etch virus;
TLS: translesion synthesis;
U2OS: human bone osteosarcoma epithelial cells;
WT: wild type;
XRCC: X-ray repair cross complementing

