## Supplemental for "Motifs of the C-terminal Domain of MCM9 Direct Localization to Sites of Mitomycin-C Damage for RAD51 Recruitment"

### Supplementary Materials

#### Supplementary Tables

**Table S1: DNA Sequences**

| DNA | Sequence (5'-3') |
|---|---|
| MCM9For*XhoI* | ATTACTCGAGCATGAATAGCGATAAG |
| MCM9648Rev*XmaI* | ATTACCCGGGCTATGACTTTTTTCTCATCTC |
| MCM9605For*XhoI* | CACCCTCGAGGGGAGGTGCACTGCTAGGAGGT |
| MCM9Rev*XmaI*Stop | ATAGATCCCGGGCTATGACTTTTTTCTCATCTCTTCATCC |
| MCM9NLS1QCF | CAACCACAGCTCCAATGCGTGTCAGTGATGATGCATCTTTTCAGCTCCGTGGGTCCACCG |
| MCM9NLS1QCR | GCAAGGGATTTAGGCCTGTCATCTTTGCCTGAGTCCAGCCTTCTGTCTGGGCTATGTACAG |
| MCM9NLS2QCF | GGGCAGACAATGTGGAAAGTAACAAGGATGATAGACTAGCACTAGATTCTGAAGCAGCAG |
| MCM9NLS2QCR | CTGCTGCTTCAGAATCTAGTGCTAGTCTATCATCCTTGTTACTTTCCACATTGTCTGCCC |
| MCM9NLS3QCF | GTTCATAGTCCTAAAATTTCCCAGGATGATGATACTAGTAGAGACGCAGCCTTGCCGGTGAAG |
| MCM9NLS3QCR | CTTCACCGGCAAGGCTGCGTCTCTACTAGTATCATCCTGGGAAATTTTAGGACTATGAAC |
| MCM9NLS4QCF | CGGTGGACCCACGGAGCTGAAAAGATGCATCATCACTGACACGCATTGGAGCTGTGGTTG |
| MCM9NLS4QCR | CAACCACAGCTCCAATGCGTGTCAGTGATGATGCATCTTTTCAGCTCCGTGGGTCCACCG |
| MCM9BRCvQCF | GGTCCAGGGGAAGAATCAAACGCCGCGACGTCATCACAGCAGGAAATCAA |
| MCM9BRCvQCR | TTGATTTCCTGCTGTGATGACGTCGCGGCGTTTGATTCTTCCCCTGGACC |
| MCM9BRCvCOF | CTGCGTAATGGTCCTGGTGAGGAAAGCAACGCAGCAACAAGCAGCCAGCAGGAGATTAACTATAGCACC |
| MCM9BRCvCOR | GGTGCTATAGTTAATCTCCTGCTGGCTGCTTGTTGCTGCGTTGCTTTCCTCACCAGGACCATTACGCAG |
| MCM9643COFor*BamHI*TEV | TTATTAGGATCCGAGAATCTATACTTTCAAGGTAGCCTGCTGAGTGAGGAGCTGCGTCGT |
| MCM9680COFor*BamHI*TEV | TTATTAGGATCCGAGAATCTATACTTTCAAGGTGGTCCTGGTGAGGAAAGCAACTTCCG |
| MCM9900CORev*XhoI* | TTATTACTCGAGGGCTAAGCTCTTAGGACGTTTGCGCTTC |
| MCM9620FNdeI | ATTACATATGCCTGAAAACCCTGGAGAGCAG |
| MCM91143RSalI | ATTAGTCGACCTATGACTTTTTTCTCATCTC |
| RAD51For*MfeI* | ATTACAATTGATGGAAAGCTTTGGCCCACAACCC |
| RAD51Rev*XhoI* | ATTACTCGAGTCTGTCTTTGGCATCTCCCACTCCATC |
| siRAD51 | GAGUUGACAAACUACUUC |

#### Supplementary Figures

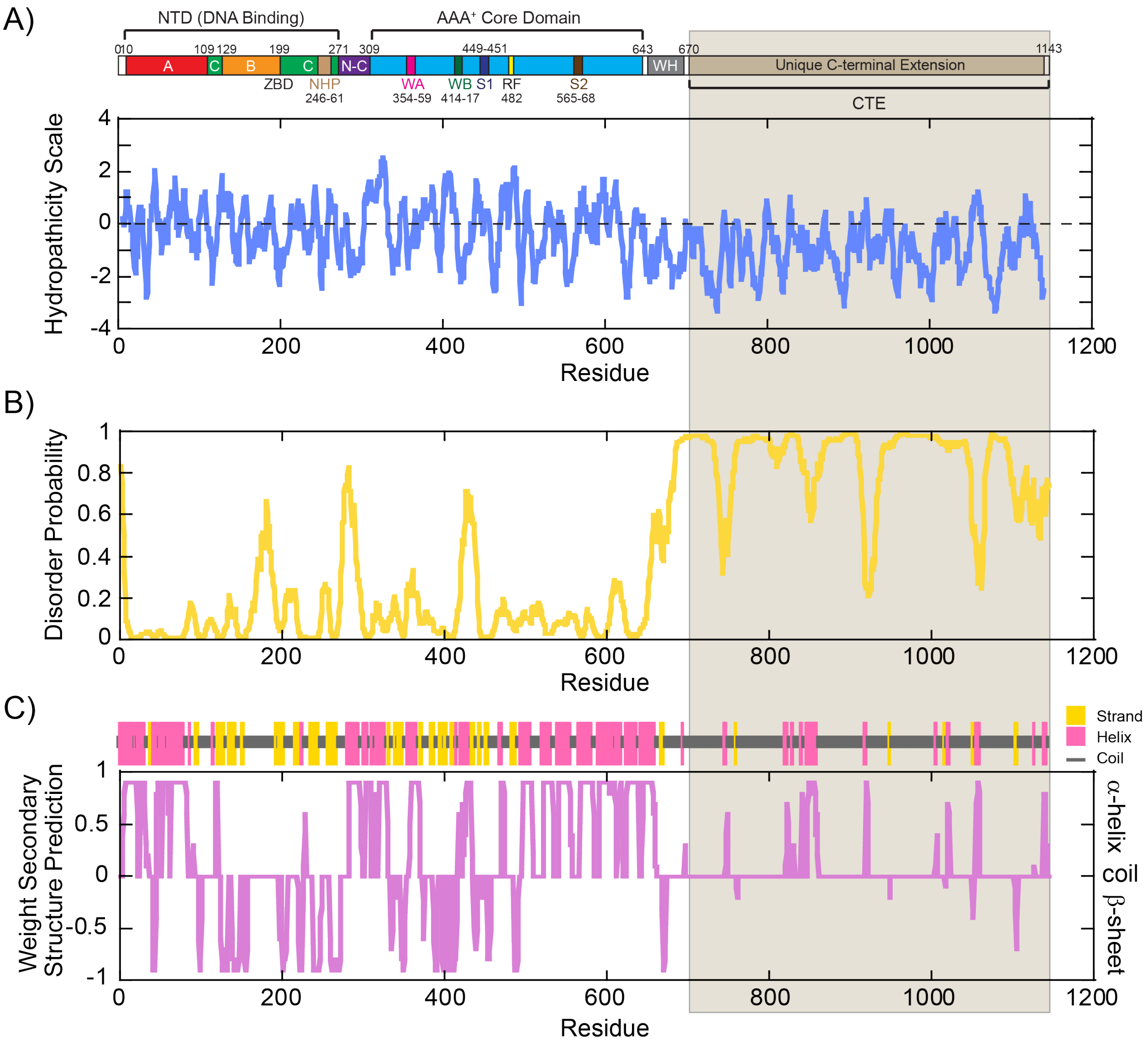


**Supplemental Figure S1. Structural prediction of the MCM9 sequence show an unstructured CTE.** The MCM9 primary protein sequence was analyzed using A) ProtScale (https://web.expasy.org/protscale/) for hydrophathicity, B) DISOPRED (http://bioinf.cs.ucl.ac.uk/disopred/) for predicted disorder, and C) PSIPRED (http://bioinf.cs.ucl.ac.uk/psipred/) for secondary structure prediction. The C-terminal half of the protein including the CTE (shaded beige) is expected to be mostly hydrophobic, disordered, and unstructured.

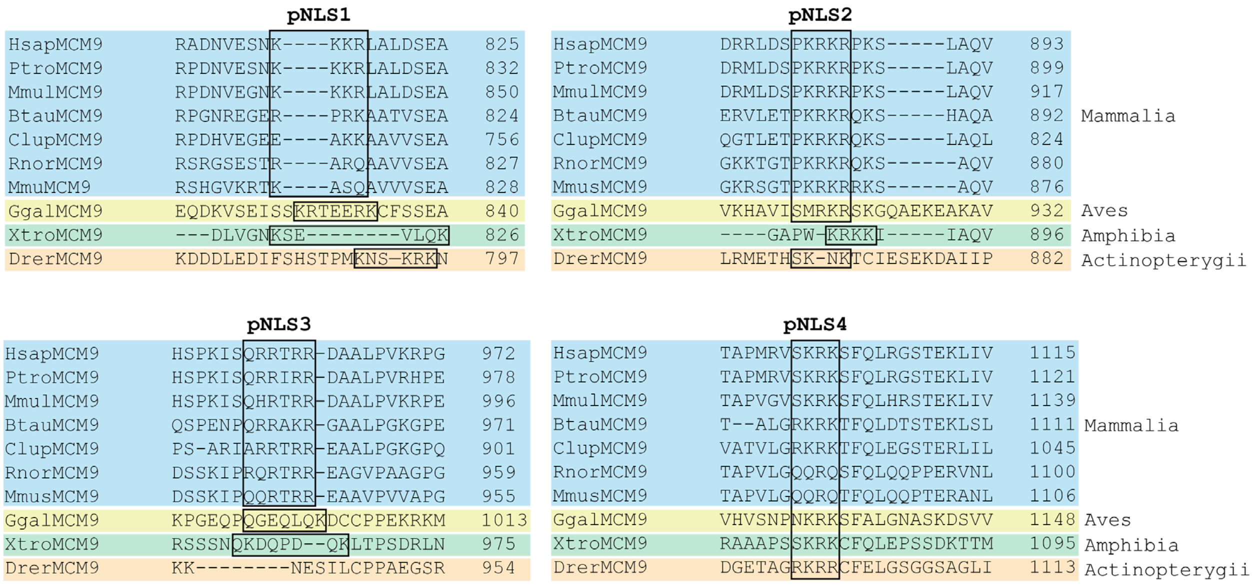


**Supplemental Figure S2.** Alignments of the NLS sequences across *Mammalia, Aves, Amphibia*, and *Actinoptergii* species. Black boxes indicate potential homology.

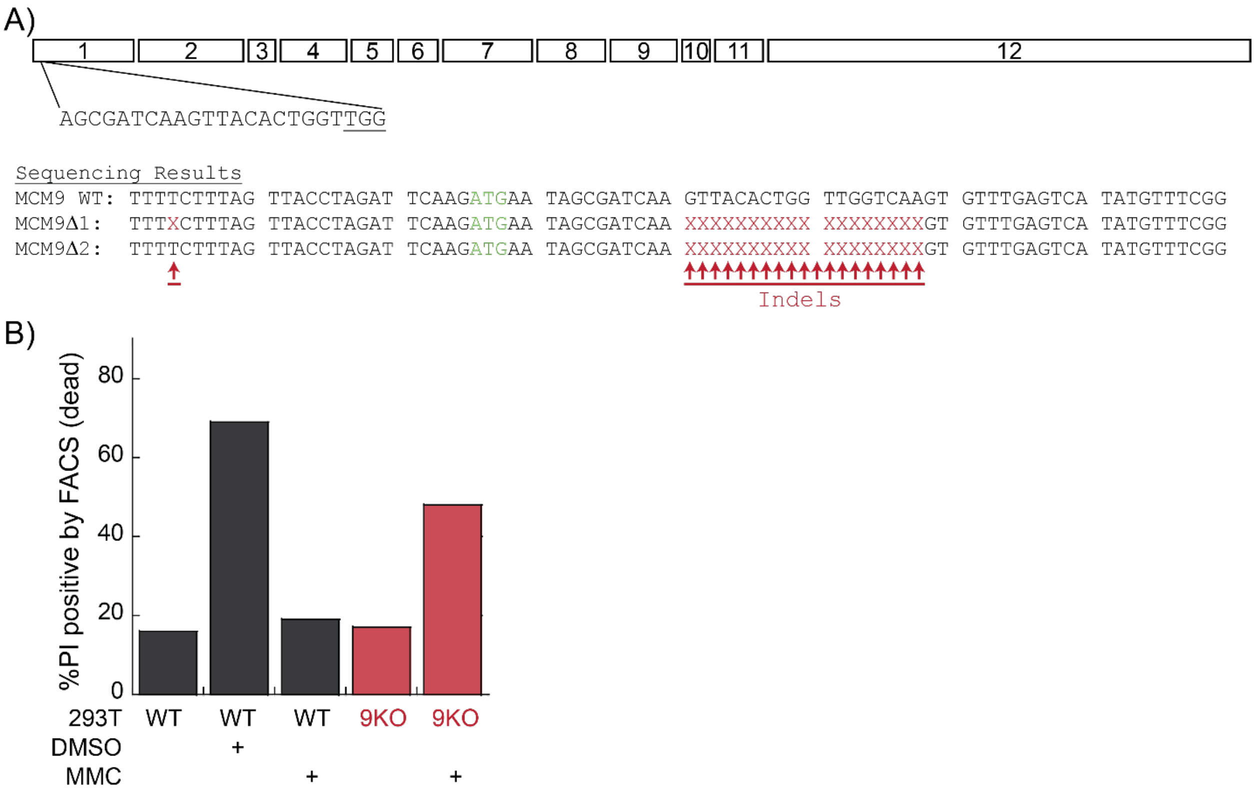


**Supplemental Figure S3. CRISPR-Cas9 knockout strategy and validation of MCM9 in HEK293T cells.** A) The 12 exons of MCM9 showing CRISPR/Cas9 targeting to exon 1. DNA Sequencing results for the selected clone deletions within exon 1 rendering a knockout of MCM9. B) Quantification of dead cells for WT 293T or 293T$^{MCM9KO}$ after treatment with MMC. DMSO was used as a positive control for cell death in WT 293T.

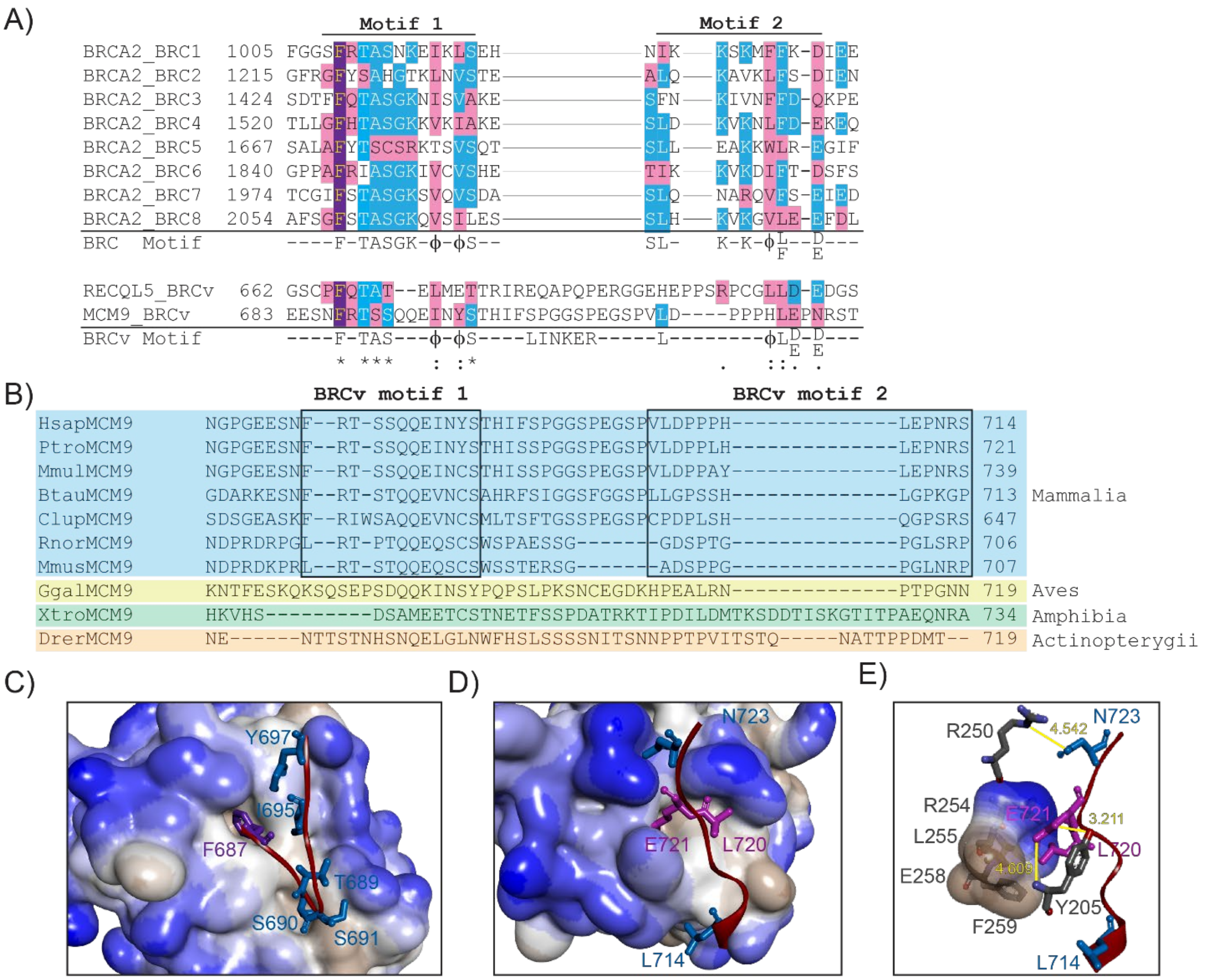


**Supplemental Figure S4. BRCv motif alignment, designation, and structure prediction.** A) Full alignment of BRC repeats of BRCA2 showing the consensus BRC motif sequence, the BRCv motif of RecQL5 and MCM9, and a consensus BRCv motif sequence. B) BRCv conservation and alignments of MCM9 across *Mammalia, Aves, Amphibia*, and *Actinoptergii* species. Closeup of C) Motif 1 and D-E) Motif 2 highlighting important interacting residues for MCM9 and RAD51 with a hydrophobic surface (white – hydrophobic and blue- hydrophilic) from a homology model in Figure 4C. Predicted H-bonds are indicated (yellow). RAD51 residues are grey.

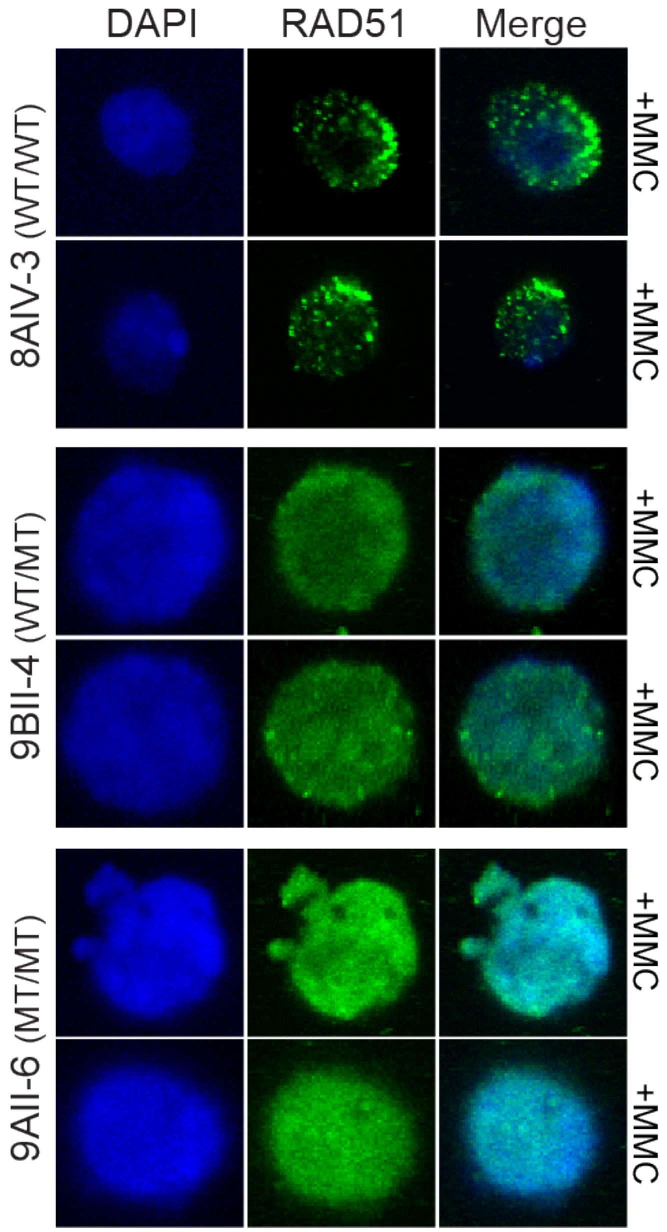


**Supplemental Figure S5. Replicates of RAD51 immunofluorescence in MCM9 patient lymphocyte cells treated with MMC.** Patient designations as in Figure 6A.
